# Estimating time-varying selection coefficients from time series data of allele frequencies

**DOI:** 10.1101/2020.11.17.387761

**Authors:** Iain Mathieson

## Abstract

Time series data of allele frequencies are a powerful resource for detecting and classifying natural and artificial selection. Ancient DNA now allows us to observe these trajectories in natural populations of long-lived species such as humans. Here, we develop a hidden Markov model to infer selection coefficients that vary over time. We show through simulations that our approach can accurately estimate both selection coefficients and the timing of changes in selection. Finally, we analyze some of the strongest signals of selection in the human genome using ancient DNA. We show that the European lactase persistence mutation was selected over the past 5,000 years with a selection coefficient of 2-2.5% in Britain, Central Europe and Iberia, but not Italy. In northern East Asia, selection at the *ADH1B* locus associated with alcohol metabolism intensified around 4,000 years ago, approximately coinciding with the introduction of rice-based agriculture. Finally, a derived allele at the *FADS* locus was selected in parallel in both Europe and East Asia, as previously hypothesized. Our approach is broadly applicable to both natural and experimental evolution data and shows how time series data can be used to resolve fine-scale details of selection.

## Introduction

Time series data of allele frequencies are obtained from many sources including experimental evolution experiments and ancient DNA studies. These data are particularly useful for estimating the strength of selection and reconstructing the allele frequencies of individual alleles. This is particularly useful when timing can be informative about the basis and environmental correlates of selection.

Many methods have been developed to solve the problem of inferring selection coefficients from time series data (Bollback *et al*., 2008; Illingworth and Mustonen, 2011; Malaspinas *et al*., 2012; Mathieson and McVean, 2013; Nishino, 2013; Feder *et al*., 2014; Lacerda and Seoighe, 2014; Foll *et al*., 2015; Terhorst *et al*., 2015; Schraiber *et al*., 2016; Ferrer-Admetlla *et al*., 2016; Shim *et al*., 2016; Nené *et al*., 2018; Paris *et al*., 2019). One assumption common to almost all these methods is that the selection coefficient is constant throughout time. This may be appropriate in some cases, for example experimental evolution where conditions are strictly controlled, but it is less appropriate in natural populations. In particular, many of the most interesting examples of human adaptation involve adaptation to new environments, gene-culture co-evolution, or infectious diseases. Selection in these cases is likely to be time-varying, and the timing of selection is typically an important question. Inferring time-varying selection requires more data than inferring constant selection, but increasing sample sizes of ancient human DNA mean that it should now be possible to infer timings and trajectories at higher resolution.

Here, we extend the hidden Markov model of Mathieson and McVean (2013) to allow selection coefficients that change over time. A model that allowed selection coefficients to vary arbitrarily would be overfitted, so we restrict selection coefficients to a pre-specified finite number of possible values and penalize changes. By defining the model in this way we are able to compute maximum likelihood estimates of the parameters using an EM algorithm.

## Methods

### Wright-Fisher model

Following the notation of Mathieson and McVean (2013), we consider a Wright-Fisher population with an effective size of 2*N_e_*. We write *f_t_* as the frequency of the selected allele at generation *t* for *t* = 0… *T*. Suppose that the frequency trajectory is known exactly and the selection coefficient *s* is constant over time. Then, an approximate maximum likelihood estimator for *s* (Watterson, 1982) is

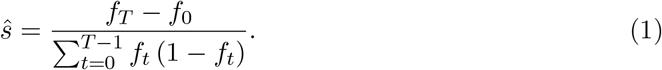

That is, the total change in allele frequency, divided by the sum of the heterozygosity over the time the allele is observed. Now suppose that the selection coefficient at generation *t* is *s_t_*, but that it takes one of *K* possible values *σ*_0_… *σ_K_*. We assume that we know which value *s_t_* takes at each generation and define indicator variables *z_t_* such that *s_t_* = *σ_z_t__*. We show in the Appendix that the maximum likelihood estimator of *σ_k_* is given by

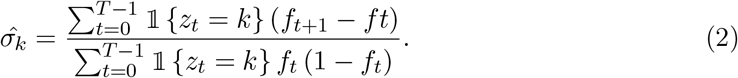

This is Equation 1 with sums over generations when the selection coefficient is equal to *σ_k_*.

### Hidden Markov model - constant selection

This model was developed in Mathieson and McVean (2013), but we describe it briefly here as background for the time-varying selection mode. In practice, *f_t_* is unknown. Instead, the data consist of samples of *n_t_* chromosomes at each generation *t* (*n_t_* can be zero), of which *a_t_* carry the selected allele. We treat *f_t_* as the hidden state in a hidden Markov model and (*a_t_, n_t_*) as the observations. To apply standard HMM theory, we discretize the frequency space so that *f_t_* ∈ *G* = {*g*_1_,…, *g_D_*}, keeping the interval between grid points *δ_g_* = *g*_*i*+1_ – *g_i_* constant. The transition probabilities **P**(*f*_*t*+1_ = *g*|*f_t_*) are computed by approximating the Wright-Fisher transition density

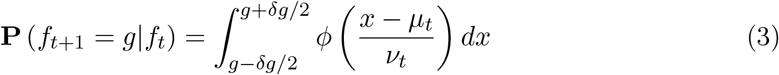

where *μ_t_* = *f_t_* + *sf_t_* (1 – *f_t_*) and 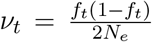. The emission probabilities are binomial *a_t_ ~ Bin* (*n_t_, f_t_*). We find the MLE for *s* by starting from an initial guess *s*^0^ and applying the EM update rule,

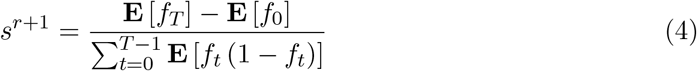

with expectations over the posterior distribution of *f_t_* computed using the forward-backward algorithm. We recalculate the forward-backward matrix and repeat until *s^r^* converges.

### Hidden Markov model - time-varying selection

In the case of time-varying selection, the hidden states are given by {*f_t_, z_t_*} for *t* = 0… *T, f_t_* ∈ {*g*_1_… *g_D_*} *z_t_* ∈ {1… *K*} The parameters are the *σ_k_* for *k* = 1… *K* (Figure 1). The emission probabilities depend only on *f_t_* and are the same as in the constant s model. The transition probabilities are given by

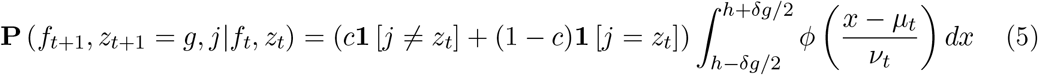

where *μ_t_* = *f_t_* + *s_t_f_t_* (1 – *f_t_*); *s_t_* = *σ_z_t__*; 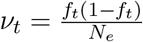 and *c* is a fixed constant that gives the probability of transitioning between hidden selection states in any generation. We show in the Appendix that the EM update rule for *σ_k_* is

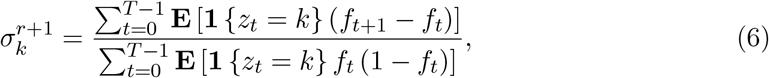

where now the expectations are taken over the joint posterior distribution of (*f_t_, z_t_*) calculated with the forward-backward algorithm. The forward-backward algorithm gives us the joint posterior probabilities 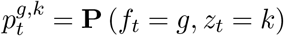, which allow us to calculate the denominator and the term **E** [**1** {*z_t_* = *k*} *f_t_*]. To calculate the term **E** [**1** {*z_t_* = *k*} *f*_*t*+1_] we also need to know the conditional posterior probabilities 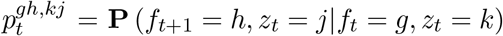 which can be computed from the forward and backward matrices. Then, Equation 6 can be written in terms of the discretized frequencies and posterior probabilities as

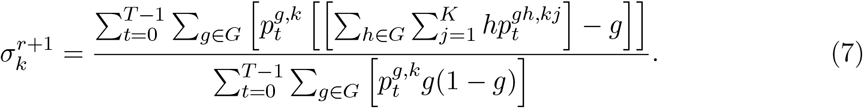

**Figure 1:**
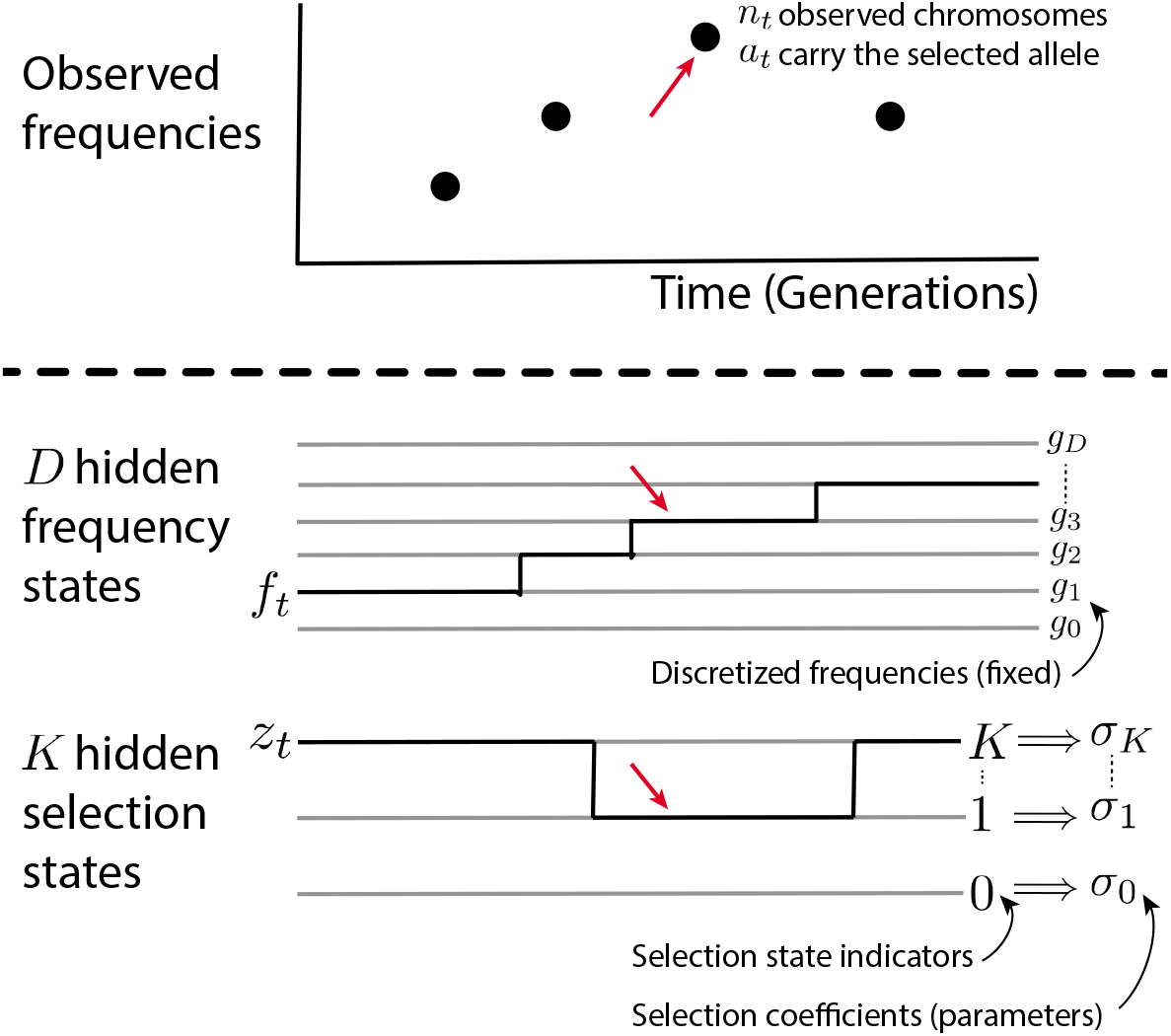
Schematic of the time-varying hidden Markov model. Below the dashed line are the hidden states. At the time indicated by the red arrows, we observe *a_t_* selected alleles out of *n_t_* total, *f_t_* = *g*_3_ and *z_t_* = 1 and therefore *s_t_* = *σ_z_t__* = σ_1_.

In summary, the algorithm is as follows:

1. Specify the number of discrete selection coefficients, *K* and the per-generation probability of changing states *c*. Make an intial guess for the selection coefficients *σ*_1_,… *σ_K_*.
2. Using the current values of *σ*_1_,… *σ_K_*, the observations *a_t_, n_t_*, the binomial emission probabilities, and the transition probabilities defined in Equation 5 compute the forward and backward matrices. Use Equation 7 to update the estimates of *σ*_1_,… *σ_K_*.
3. Repeat step 2 until iteration *r* where 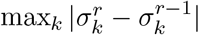 is less than some pre-defined tolerance, and stop.

Because there are *DK* hidden states, running time is *O*(*D*^2^*K*^2^*T*) and space is *O*(*DKT*).

### Simulated data

We simulated allele frequencies under a Wright-Fisher model, with an effective population size of *N_e_* = 10,000 under three different scenarios (Fig. 2A-C);

1. The selection coefficient is 0.02 for 50 generations and then –0.02 for 50 generations. Initial frequency *f*_0_ = 0.1.
2. The selection coefficient is 0.02 for 100 generations, 0 for 50 generations, and then –0.02 for 50 generations. Initial frequency *f*_0_ = 0.1.
3. The selection coefficient alternates between 0.02 and –0.02 every 40 generations. Initial frequency *f*_0_ = 0.5.

**Figure 2:**
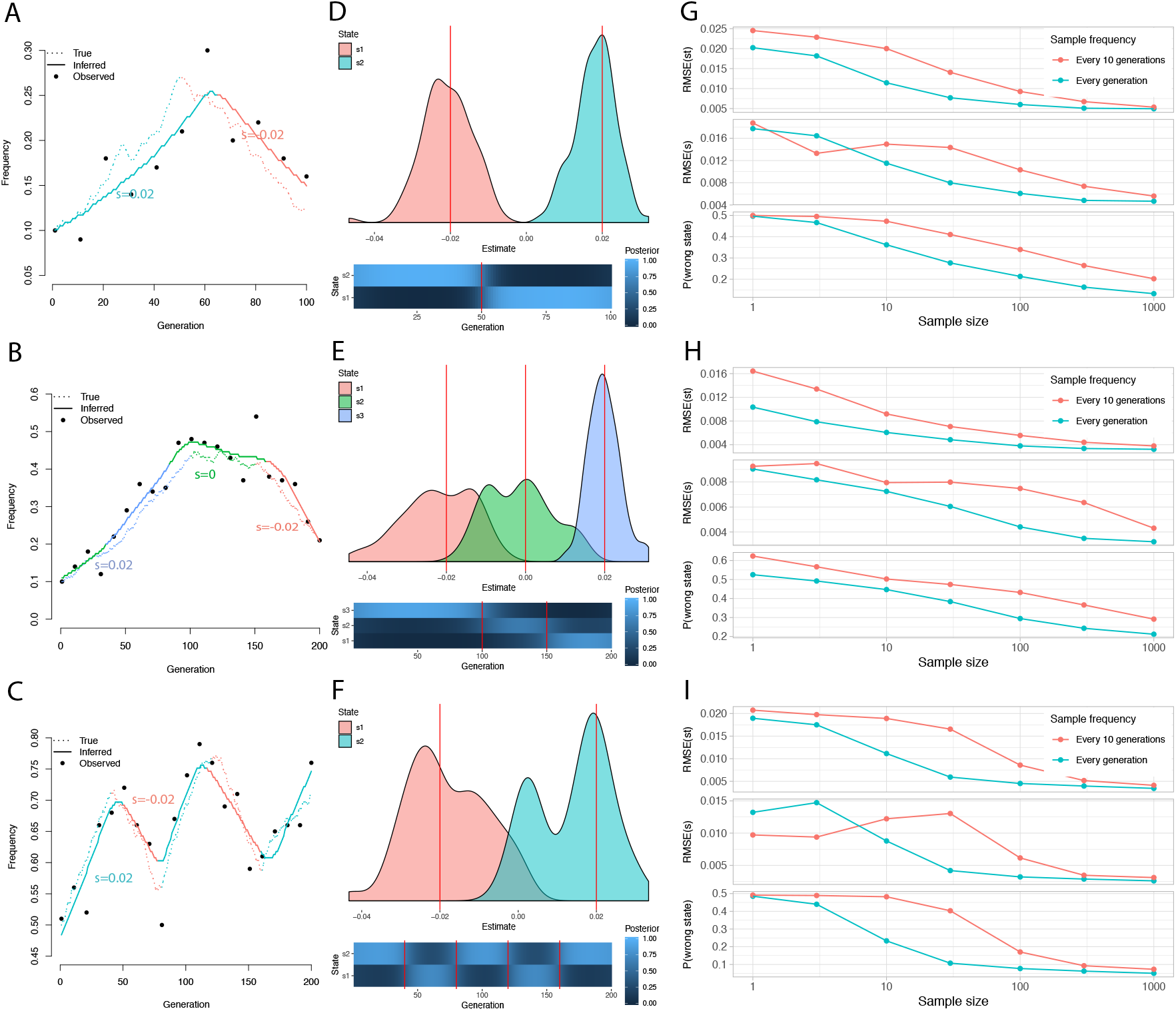
Performance of the estimator on simulated data. **A-C**: Simulated trajectories (dashed), observations (points), and inferred trajectories (solid). Colors indicate true and inferred selection states. **D-F**: For each of the scenarios in A-C, density plots of distribution of the estimates of the selection coefficients 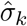 from 100 simulations. Red lines mark the true values. Lower panels show the average posterior probabilities of being in each selection state (**P**(*z_t_* = *k*)) in each generation. Red lines mark the true changepoints. **G-I**: For each of the scenarios in A-C, we show the RMSE error in 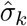 in the upper panel, the RMSE error in *ŝ_t_* in the middle panel, and the posterior probability that *z_t_* is wrong in ±**10** generations around each changepoint. We show estimates for sample sizes ranging from 1 to 1000, sampled either every generation or every 10 generations.

We sampled 100 haploid individuals every 10 generations. We set initial estimates of *σ_k_* to be ±0.05 for *K* = 2 and 0, ±0.05 for *K* = 3, a grid size of 100 (i.e. *D* = 100) and a tolerance of 0.001. We fixed the probability of transitioning between selection states c to be the inverse of the total generations observed; i.e. we expect ~ 1 selection state transition. We show the distribution of the point estimates of 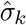, and the averaged posterior distribution of the *z_t_* (Fig. 2D-F). Finally, we varied both the frequency and size of the samples and investigated how the performance of the estimator changed (Fig. 2G-I) in terms of:

- The root mean squared error in the estimate of the selection coefficients 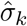.
- The posterior probability that the the inferred selection state is correct within ±10 generations of each changepoint.
- The root mean squared error in the weighted per-generation estimate of the selection coefficient 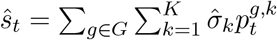.

We investigated performance as we varied parameter values and specified incorrect values for fixed parameters, for example *N_e_, c* or *K*.

### Comparison with existing approaches

We compared our approach to CP-WFABC (Shim *et al*., 2016)—the only existing method that is able to infer time-varying selection coefficients. Specifically, CP-WFABC uses Approximate Bayesian Computation (ABC) to fit a model with a single changepoint and two selection coefficients (i.e our scenario 1). We used the default number of simulations (1,000,000) with the best 1,000 retained, and set the prior to be the range (–2*s*, 2*s*) as we tested performance for different values of s. We use the posterior distribution of the change-point to calculate the probability of being in the wrong state, and the posterior mode as a point estimate of the selection coefficients which we compare with our maximum likelihood estimates.

### Ancient DNA data

We collected published ancient DNA data from four regions of Europe chosen because they had large sample sizes and corresponding present-day data from the 1000 Genomes Project (1000 Genomes Project Consortium, 2015). We restricted to dates after the arrival of Steppe-related ancestry in each region to minimize the effects of changes in ancestry associated with that arrival (Haak *et al*., 2015). The four regions were: Britain (GBR, 50-60°N, 5°W-2°E, <4400BP), Central Europe (CEU, 47-53°N, 8-20°E, <5000BP), Italy (TSI, 36-45°N, 7-15°E, <5000BP), Iberia (IBS, 36-44°N, 10°W-4°E, <5000BP). We identified a total of 499 samples, although not all had coverage at rs4988235 or rs174546. The samples were originally published in the following references: Allentoft *et al*. (2015); Amorim *et al*. (2018); Antonio *et al*. (2019); Fernandes *et al*. (2018); Gamba *et al*. (2014); Lipson *et al*. (2017); Martiniano *et al*. (2016, 2017); Mathieson *et al*. (2015, 2018); Mittnik *et al*. (2019); Narasimhan *et al*. (2019); Olalde *et al*. (2018, 2019); Schiffels *et al*. (2016); Valdiosera *et al*. (2018); Veeramah *et al*. (2018) and Zalloua *et al*. (2018). We also identified 255 ancient samples from East Asia (excluding Japan) from Ning *et al*. (2020); Yang *et al*. (2020) and Wang *et al*. (2020) and divided them into “North” and “South” populations at 30°N. We restricted the South population to <5000BP because only one sample was older.

### Ancient DNA analysis

We used a grid of *D* = 1000, two selection states and a tolerance of 1 × 10^-4^. We set *N_e_* to grow exponentially from 10^4^ to 10^6^ over the past 200 years approximately as inferred by Browning and Browning (2015), though without the more rapid increase in past 10 generations. Though this estimate is for European populations, our estimator is robust to mis-specification of *N_e_* so we assumed it was representative of late Holocene growth rates and used the same values for East Asia. Finally, we estimated the bias and uncertainty in our estimates using a parametric bootstrap: we simulated observations conditional on the inferred frequency trajectory and actual sample dates, and then reran the estimator.

### Logistic regression analysis

We ran an independent analysis where we fitted the observations using logistic regression on time and ancestry components estimated using ADMIXTURE with K=3 (Alexander *et al*., 2009). That is, the expected allele frequency of individual *i, f^i^* is given by:

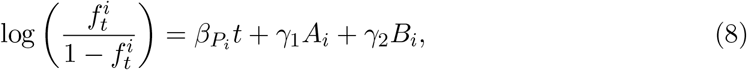

where *P_i_* is the population to which individual *i* belongs and *A_i_* and *B_i_* are two of its ancestry component values (the third is 1 – *A_i_* – *B_i_*). We estimate s by estimating the predicted change in frequency in one generation for each individual, converting it to an estimate of *s* based on the expected frequency change in the Wright-Fisher model (i.e. 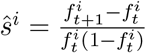) and then averaging over all individuals in each population. We estimate the standard error by assuming that the ratio of *ŝ* to its standard error is the same as the ratio of *β_P_i__* to its standard error. While this is not an explicit model of the evolutionary process, it does allow us to account for variation in genome-wide ancestry across individuals.

## Results

### Simulated data

In simulated data, we recover allele frequency trajectories, selection coefficients and the timing of changes in selection coefficients (Fig. 2). Simulations also allow us to test the robustness of the estimator to misspecification and highlight key features of its behavior. First, under scenario 1, we tested robustness to misspecification of *N_e_* and *c*. These parameters must be specified in advance. However, we find that the error in the estimates is robust over one order of magnitude for *N_e_*, and two orders of magnitude for *c* (Fig. S1 & S2) Thus, as long as reasonable estimates of these parameters are available, misspecification should not be a major concern. Second, we note that even for very large samples the RMSE of the selection coefficient 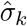 and *ŝ_t_* do not tend to zero. This is partly due to the stochastic effect of drift and partly due to the fact that the estimators can be biased, particularly for low initial frequencies (Fig. S3). If the initial frequency is very low, there is a relatively high chance that the allele is just by drift. For example, for an allele in a single copy, there is a probability of ~ e^-1^ ≈ 0.37 that the allele is lost in one generation leading to a negative MLE for the selection coefficient.

As sample size increases, the RMSE of *ŝ_k_* decreases more reliably than that of 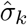 (Fig. 2G-I). In other words, the estimator is better at answering the question “what is the selection coefficient in generation *t*?” than “what is the selection coefficient in state *k*?”. The first question allows us to average estimates over multiple states, even if the number of states is misspecified. In fact, if there are too many or too few selection states in the HMM, then the estimator does over- or underestimate the number of transitions (Fig. S4A) but the error in *ŝ_t_* does not change (Fig. S4B). Therefore in our analysis of real data we focus on *ŝ_t_*, rather than 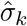.

In practice, the performance of the estimator depends on the data. For example, the accuracy with which we are able to detect fluctuating selection in scenario 3 (Fig. 2C) depends on the period of fluctuation (Fig. S5). Performance also depends on the sampling scheme. If we do not sample around a changepoint then we will misestimate selection coefficients around that time. Given relatively smooth trajectories, performance depends on the total number of observations—sampling ten times as many chromosomes ten times less frequently gives about the same error (Fig. 2G-I). However more uniform sampling in time would be more robust to rapidly changing trajectories. In general we recommend assessing the performance and robustness of the estimator using a parametric bootstrap approach. Run the estimator on the observed data, simulate data under the inferred model and actual pattern of observations, and investigate performance on the simulated data.

Finally, we compared the performance of our estimator to the only previously published method for detecting time-varying selection coefficients—CPWFABC (Shim *et al*., 2016). This method uses Approximate Bayesian Computation to jointly infer a single changepoint and two selection coefficients (pre- and post-changepoint). We tested the performance of this model under scenario 1 and find that our estimator outperforms it both in terms of locating the changepoint and estimating the selection coefficients (Fig. S6).

### Selection at *LCT* in Europe

The SNP rs4988235 (C/T-13910) is associated with adult lactase persistence in Europeans (Enattah *et al*., 2002) and exhibits one of the strongest signals of positive selection in the entire genome (Bersaglieri *et al*., 2004; Grossman *et al*., 2013). Estimates of the strength and timing of selection on the variant based on present-day data are variable and have wide confidence intervals, ranging from 0-0.2 for *s*) and ~1500-65,000 years before present for the origin of the mutation (Bersaglieri *et al*., 2004; Tishkoff *et al*., 2007; Itan *et al*., 2009; Peter *et al*., 2012). Direct evidence from ancient DNA has established that the allele was rare or absent in the Neolithic and was not present at substantial frequency until the Bronze Age, starting around 5000BP (Burger *et al*., 2007; Allentoft *et al*., 2015; Mathieson *et al*., 2015). In parts of Europe, for example Iberia, the derived allele did not become common until even later (Olalde *et al*., 2019). Using ancient DNA data from across Europe, Mathieson and Mathieson (2018) estimated a selection coefficient of 0.018.

We used data from 499 ancient Europeans, divided by region, to investigate whether there were differences in the selective pressure across Europe, and whether the strength of selection varied over time (Fig. 3). We estimate that in Britain and Central Europe, the variant experienced a selection coefficient of ~0.025, consistently for the past 4-5000 years. In Iberia, the selection coefficient was slightly lower—around 0.02. Bootstrapping suggests that the selection coefficients outside Italy might be underestimated by up to 0.005 (Fig. 3). We find no evidence that the allele was ever under selection in Italy, with an estimated selection coefficient of zero. One concern is that these differences might be due to difference in the timing of ancestry changes across Europe. We therefore fitted a logistic regression to the observations, including date and two ancestry components (inferred using ADMIXTURE with *K* = 3). This model yields similar estimates of the selection coefficients (Fig. S7). Finally, we fitted the lattice model from Mathieson and McVean (2013) allowing migration between demes and, again, find very similar results (Fig. S8).

**Figure 3:**
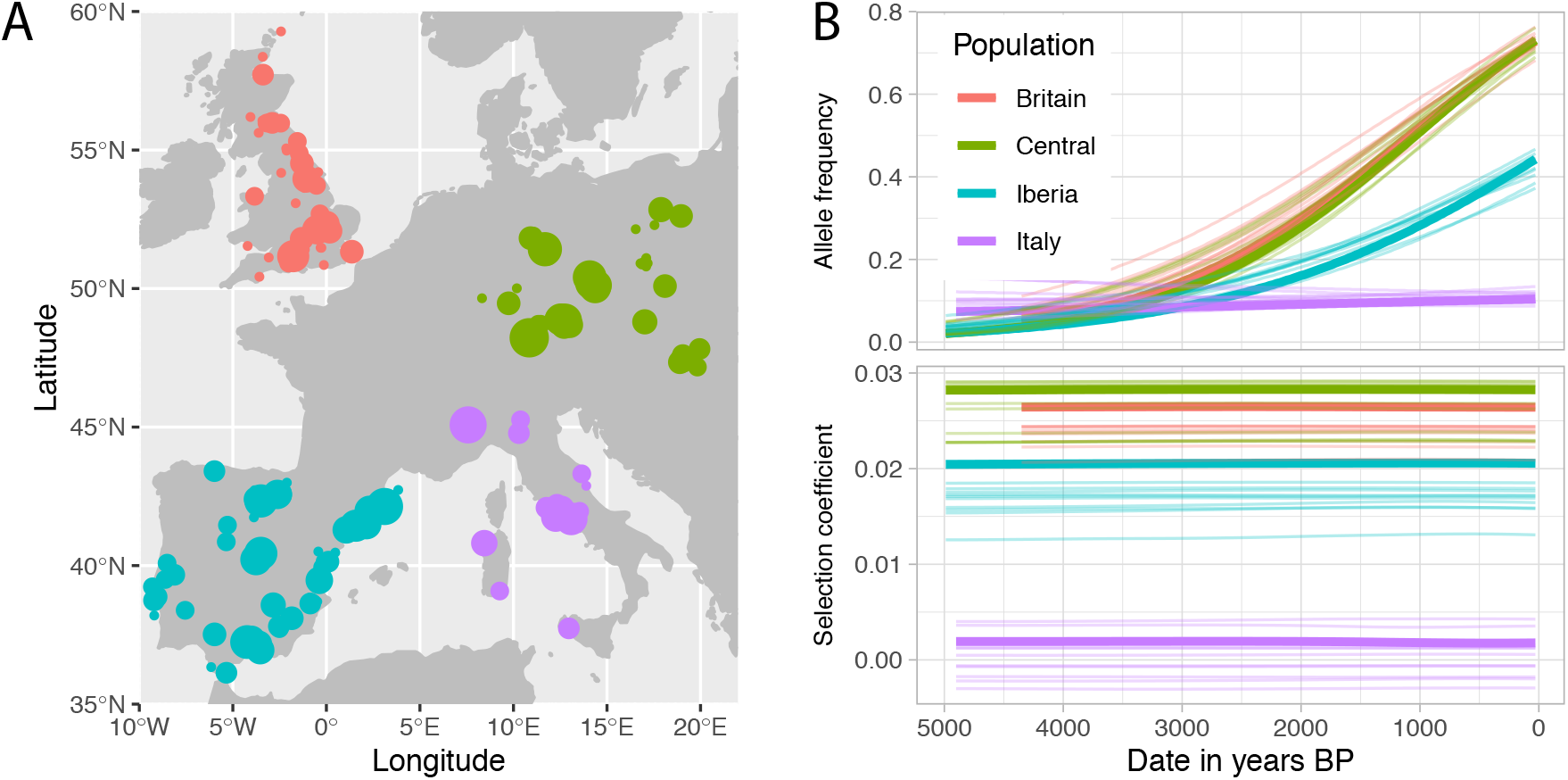
Selection at *LCT*. **A**: Location of 499 samples used in the analysis. The area of each circle is proportional to the sample size at each site. **B**: **Upper panel**: Solid lines indicate the inferred allele frequency trajectory for the lactase persistence allele in different parts of Europe. Faded lines indicate bootstrap replicates generated by sampling observations from this inferred frequency trajectory **Lower panel**: Inferred selection coefficient (*ŝ_t_*) and bootstrap replicates as a function of time.

It is unknown whether selection on lactase persistence was dominant or additive. If we assume that the selection coefficient is constant over time, we can test the effect of different dominance parameters (Mathieson and McVean, 2013). Maximum likelihood estimates indicate complete or partial dominance, but the difference in log-likelihood is small and we cannot reject additivity (Fig. S9). Finally, it has been suggested that the allele had already reached its present-day frequency by the Middle Ages (Kruttli *et al*., 2014) and that selection must have stopped by then. Simulations show that, given the distribution of observations, we would be unable to detect this change in selection, so this question remains unresolved (Fig. S10).

### Selection at *ADH1B* in East Asia

The alcohol and aldehyde dehydrogenase genes *ADH1B* and *ALDH2* are the key components of the oxidative alcohol metabolism pathway. The derived A allele of rs1229984 in *ADH1B* increases the rate at which ethanol is oxidised to acetaldehyde and the A allele of rs671 in *ALDH2* decreases the rate at which acetaldehyde is transformed into acetic acid. The net effect of the two polymorphisms is to increase the concentration of acetaldehyde after consuming alcohol, leading to unpleasant negative effects; consequently the variants are protective against alcohol abuse (Chen *et al*., 1999). These two variants are at high frequency in East Asia (0.8 and 0.2, respectively) compared to the rest of the world (up to 0.03 and 0.00) (1000 Genomes Project Consortium, 2015). Both variants exhibit genomic signatures of selection (Oota *et al*., 2004; Barreiro *et al*., 2008; Okada *et al*., 2018). Explanations include protection against alcohol abuse and the anti-parasitic action of aldehyde (Oota *et al*., 2004), and the variants are thought to be associated with the Neolithic development of rice farming (Peng *et al*., 2010). Using ancient DNA from 255 ancient individuals from East Asia (Ning *et al*., 2020; Yang *et al*., 2020; Wang *et al*., 2020), and present-day allele frequencies from 1103 individuals (Peng *et al*., 2010), we estimated the frequency and selection coefficient trajectories for *ADH1B* (Fig. 4). We estimate that by 4000 BP, the derived *ADH1B* was already common south of 30°N, but was still rare further north. Selection intensified in the north around 4000 BP with a selection coefficient of around 2%. We find consistent results if we replace the present-day population samples with the CHB and CHS 1000 Genomes populations, and when we fit the logistic regression model, correcting for K = 3 inferred ancestry components (Fig. S11).

**Figure 4:**
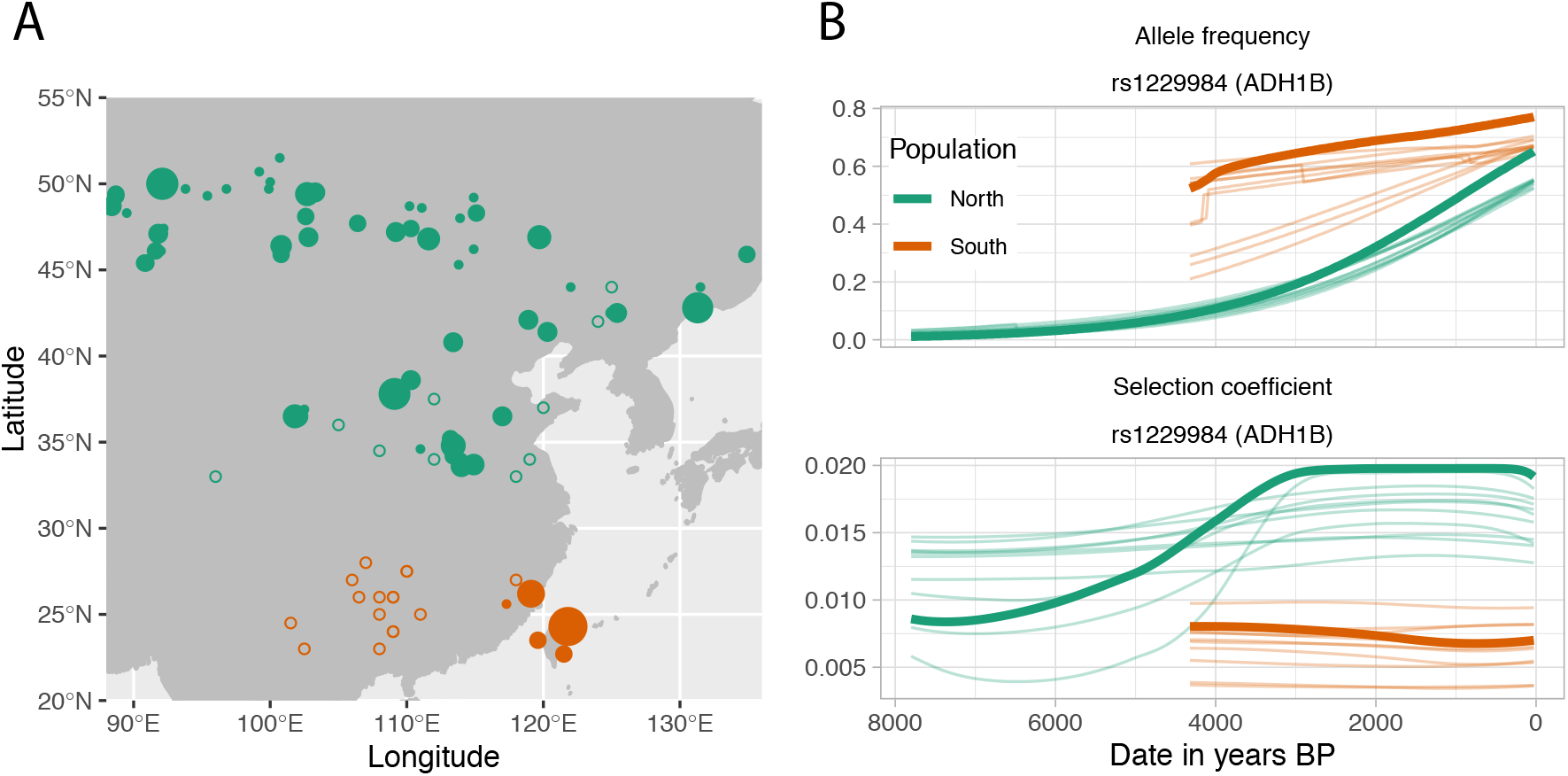
Selection at *ADH1B*. **A**: Location of samples used in the analysis. The area of each circle is proportional to the sample size at each site. Open circles denote locations of present-day samples. **B**: **Upper panel**: Solid lines indicate the inferred allele frequency trajectory for the derived *ADH1B* allele in North and South East Asia. Faded lines indicate bootstrap replicates generated by sampling observations from the inferred trajectory **Lower panel**: Inferred selection coefficient (*ŝ_t_*) and bootstrap replicates as a function of time.

Rice was domesticated in the Yangtze basin (≈ 30°N) as early as 8000 BP and our results suggest that by 4000 BP, the derived *ADH1B* allele was common there. It subsequently spread north where it experienced strong selection. We did not find the derived *ALDH2* allele in any ancient individuals suggesting that it was selected in both north and south East Asia in the past few thousand years on a background of the derived ADH1B allele.

### Selection at *FADS* in Europe and East Asia

Another signal of selection in Europe is found at the *FADS* locus. Here the derived variant has been strongly selected in the past 10,000 years and is thought to be an adaptation to an agricultural diet (Ameur *et al*., 2012; Mathieson *et al*., 2015; Buckley *et al*., 2017; Ye *et al*., 2017; Mathieson and Mathieson, 2018) In contrast to the LCT locus, we find that the derived allele at the FADS locus tagged by rs174546 follows approximately the same trajectory in each region, and has approximately the same selection coefficient (0.007-0.012), consistent with a Europe-wide estimate of 0.004-0.015 (Mathieson and Mathieson, 2018) (Fig. 5A). In East Asia, we find that the same allele has also been under recent selection, with a trajectory and selection coefficient in the north that is similar to that observed in Europe (Fig. 5B). In the south we estimate a lower frequency but stronger selection though with only one observation (out of 30) of the derived allele, this is very uncertain. In both cases, we find consistent results with the logistic regression model (Figures S12 and S13).

**Figure 5:**
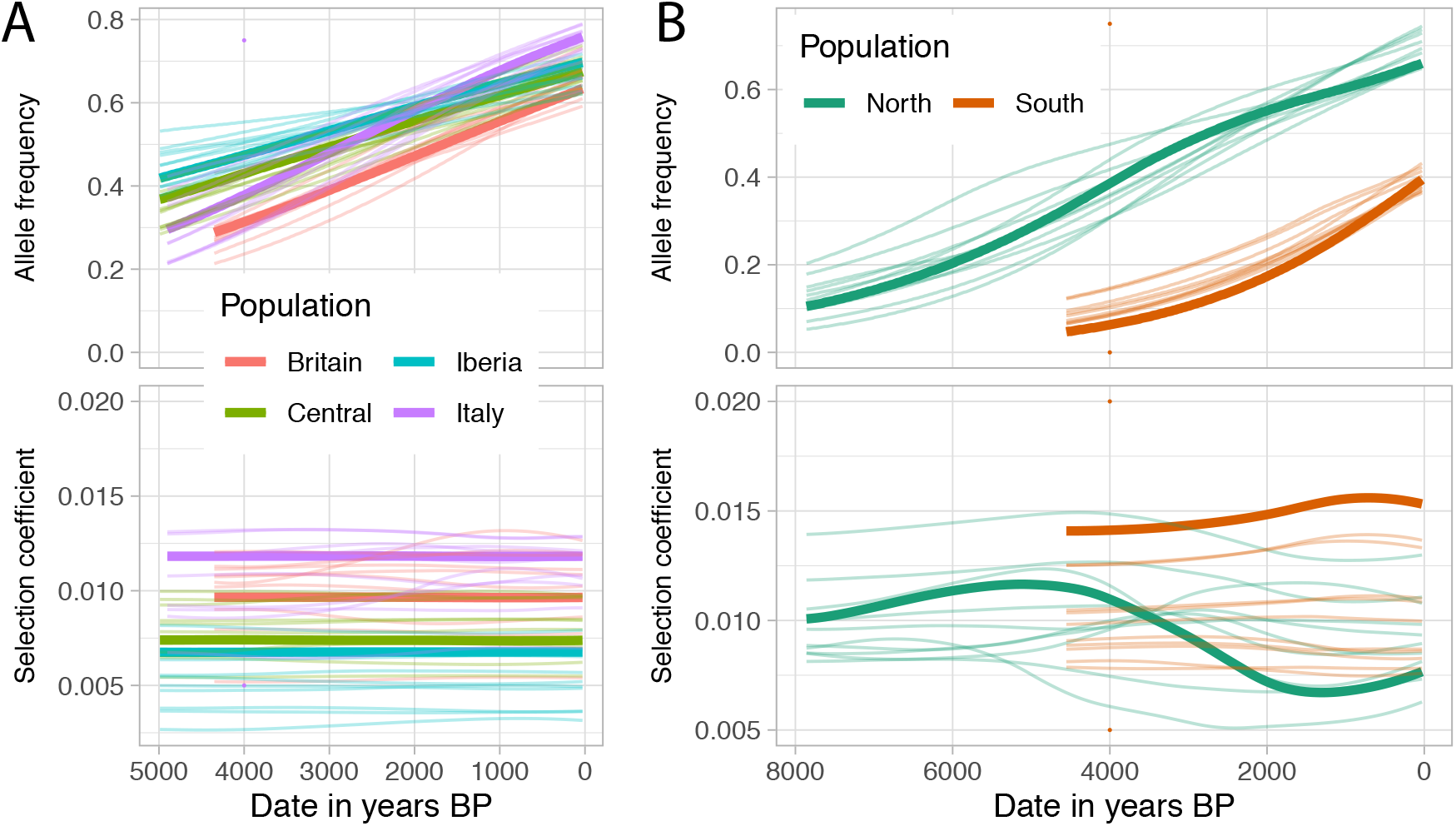
Inferred allele frequency trajectories and selection coefficient for the derived *FADS* allele in **A** Europe and **B** East Asia. Details are as in Figures 3 and 4. Present-day allele frequencies taken from the 1000 Genomes project populations.

## Discussion

Ancient DNA is powerful tool for studying the role of natural selection in human evolution. By detecting time-varying selection, we can identify environmental changes leading to selective pressure on particular alleles. Our approach is not limited to human data, and is broadly applicable to ancient DNA, ecological or experimental evolution studies.

We find that the selection coefficient for the European lactase persistence allele was consistently around 2-2.5% in Britain, Central Europe and Iberia while the allele was not selected at all in Italy. The distribution of observations mean that we have limited power to detect changes in selection coefficient over this time period. In East Asia, our analysis of the ADH1B locus is consistent with selection intensifying in the North after 4000 BP, corresponding to the introduction of rice farming. However, geographic sampling and knowledge of ancestry changes is currently more limited in East Asia than in Europe, so this result does not exclude more complex trends. As previously hypothesized (Mathieson, 2020), the derived FADS allele was selected in both Europe and East Asia.

Genomic signatures of selection are relatively easy to detect with present-day data. Ancient DNA provides temporal information, as well as information about changes in ancestry, allowing the timing and strength of selection to be inferred. Though this does not solve the ultimate problem of identifying the environmental drivers of selection, it goes a long way to making that problem tractable, allowing hypotheses to be rejected. For example, one hypothesis about selection for lactase persistence is that it allows the uptake of vitamin D from milk rather than UV radiation, which is advantageous in the North but not South of Europe. However, our results show that selection was almost as strong in Iberia as in Northern Europe and much stronger than in Italy, making this unlikely to be the sole explanation. By allowing these inferences, our approach and others based on ancient DNA should provide much deeper insight into the nature of recent human evolution.

## Acknowledgments

We thank Ziyue Gao for helpful comments on an earlier version of the manuscript. This research was funded by grants from the Alfred P. Sloan Foundation [FG-2018-10647], the Charles E. Kaufman Foundation [KA2018-98559], and NIGMS [R35GM133708]. The content is solely the responsibility of the author and does not necessarily represent the official views of the National Institutes of Health or other funding sources.

## Data availability

An R package is available at https://github.com/mathii/slattice/

## Appendix

This derivations follow very closely those for the constant selection case in Mathieson and McVean (2013). Suppose the allele frequency *f_t_* in generation *t* is known exactly. The selection coefficient in generation *t* is 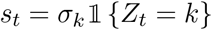 where *z_t_* is known. Then, conditional on *f_t_*, the distribution of *f*_*t*+1_ is binomial with size *N_e_* and probability *f_t_* + *s_t_f_t_*(**1** – *f_t_*). Thus, log-likelihood of the selection coefficients *σ*_1_… *σ_K_* is given by:

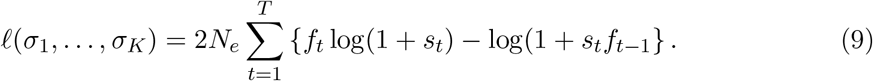

But, since 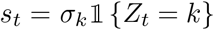, the log-likelihoods for each *σ_k_* do not depend on each other so we can write

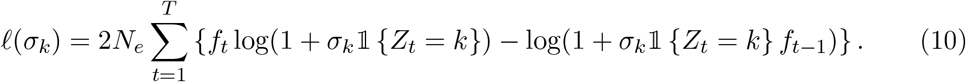

Differentiating w.r.t. *σ_k_* and setting equal to zero gives.

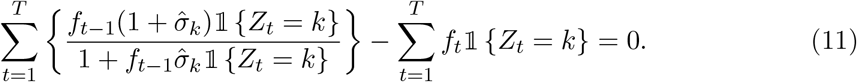

Expanding the fraction to first order in *σ_k_* gives

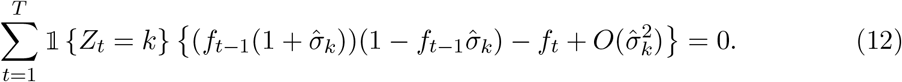

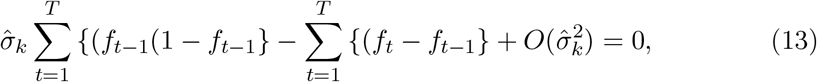

which yields the result in Equation 2. Another way to see this is that in Equation 10, we could remove the indicator functions and write the sum over *t*: *Z_t_* = *k*, rather than *t* = 1… *T* leading to an equivalent form of Equation 2. For the EM update step we maximize the expectation over {*f_t_, z_t_*} of the likelihood (Equation 10). Taking expectations, differentiating and setting equal to zero we obtain, by the same argument above, the result of Equation 6.

## Supplementary Figures

**Figure S1:**
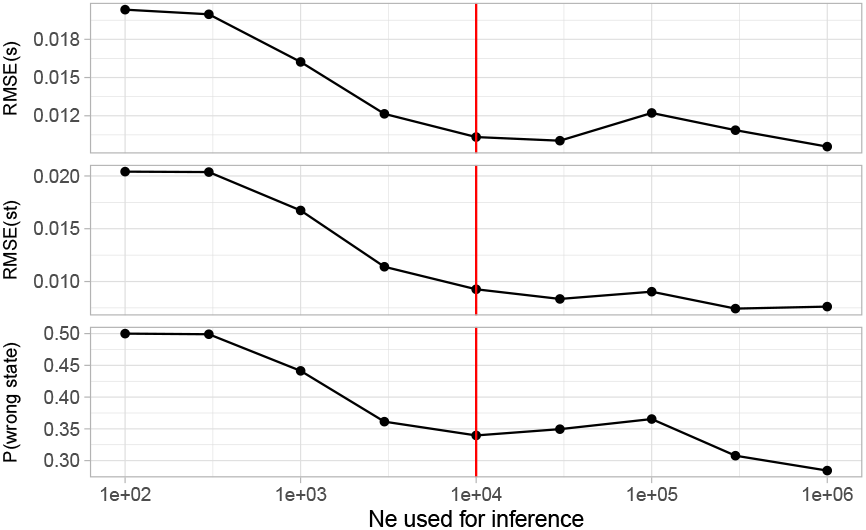
Errors in scenario 1 (defined as in Fig. 2G) when *N_e_* is mis-specified. True *N_e_* = 10, 000, and we sample 100 chromosomes every 10 generations.

**Figure S2:**
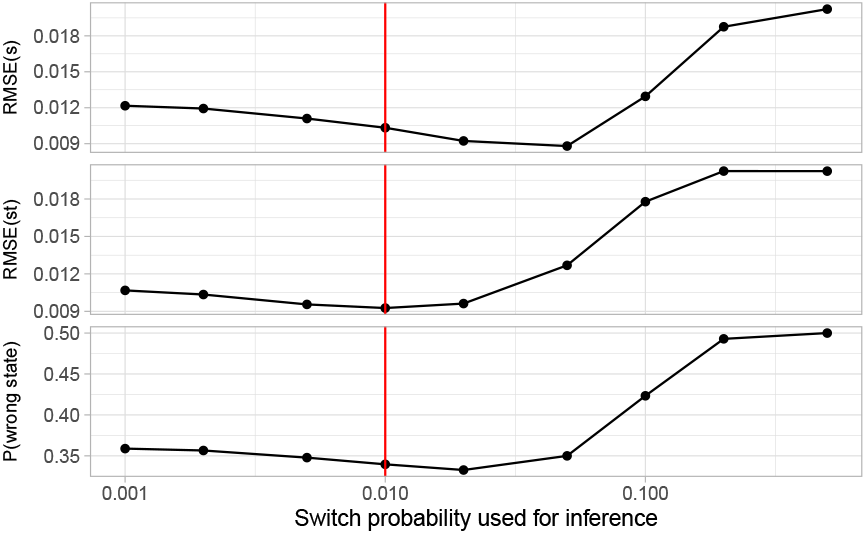
Errors in scenario 1 (defined as in Fig. 2G) when *c* is mis-specified. True *c* = 0.01, and we sample 100 chromosomes every 10 generations.

**Figure S3:**
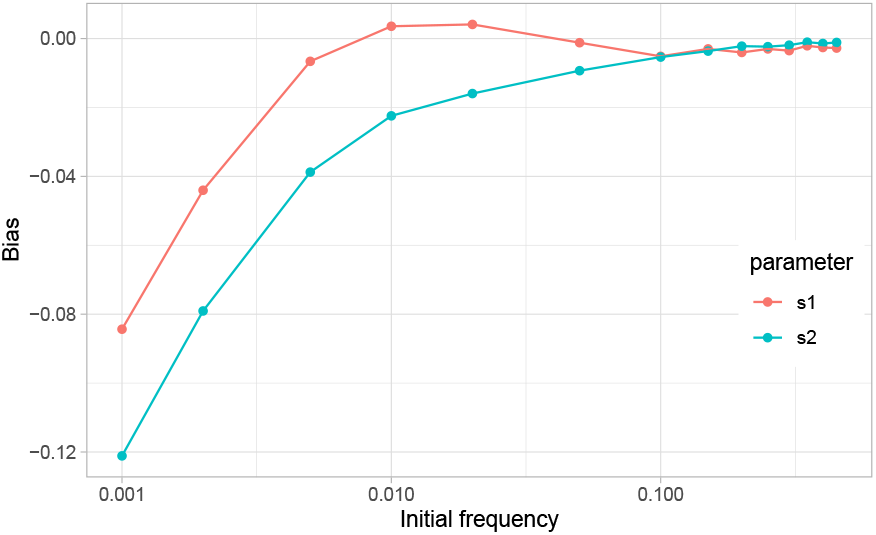
Bias in the estimate of selection coefficients 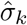 in scenario 1 as a function of initial allele frequency. Simulations as in Fig. 2D.

**Figure S4:**
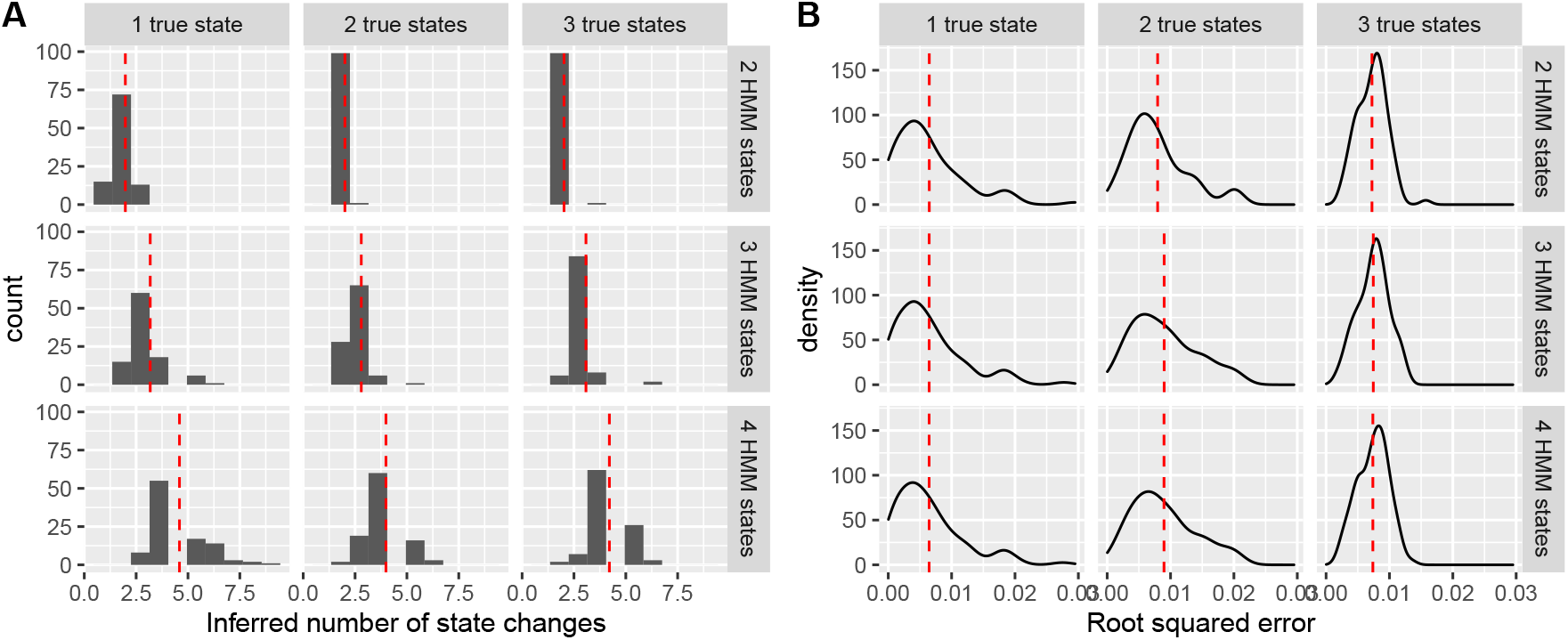
Performance of the estimator when the number of selection states is misspecified. **A**: distribution of the number of inferred state changes (in the sense that the most likely state changes), for different numbers of true model states. Histograms show the distribution of inferred state changes from 100 replicates, and dashed red lines show the mean. For 1 true state we simulate *s* = 0.02 for 50 generations, for 2 states we simulate *s* = 0.02 and 0 for 50 generations each, and for 3 states we simulate *s* = 0.02, 0 and −0.02 for 50 generations each. **B**: With the same simulations as part A, we show the distribution of RMSE of *ŝ_t_* for different numbers of model states. Dashed red lines show the mean.

**Figure S5:**
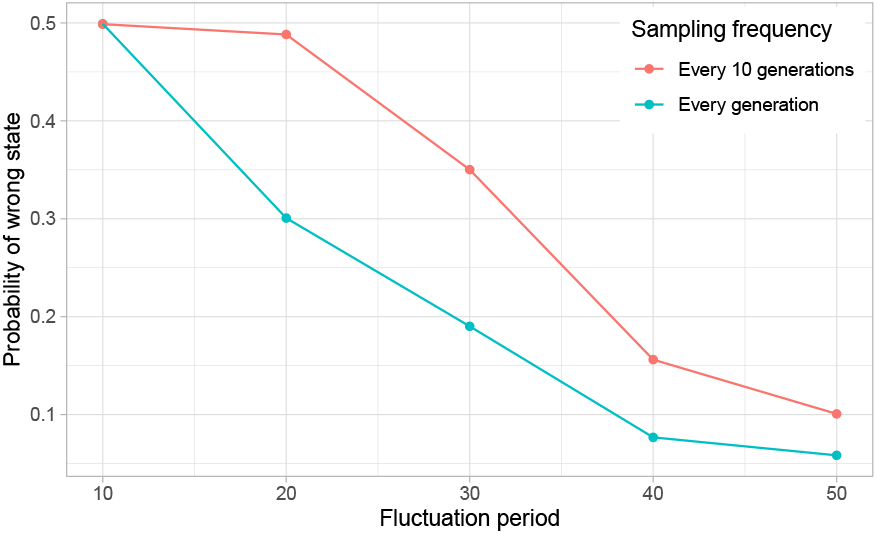
Performance of the estimator for scenario 3 (Fig. 2C) when the period of fluctuation varies. We show the probability that we estimate that we are in the wrong state. Observations are 100 chromosomes either every generation or every 10 generations.

**Figure S6:**
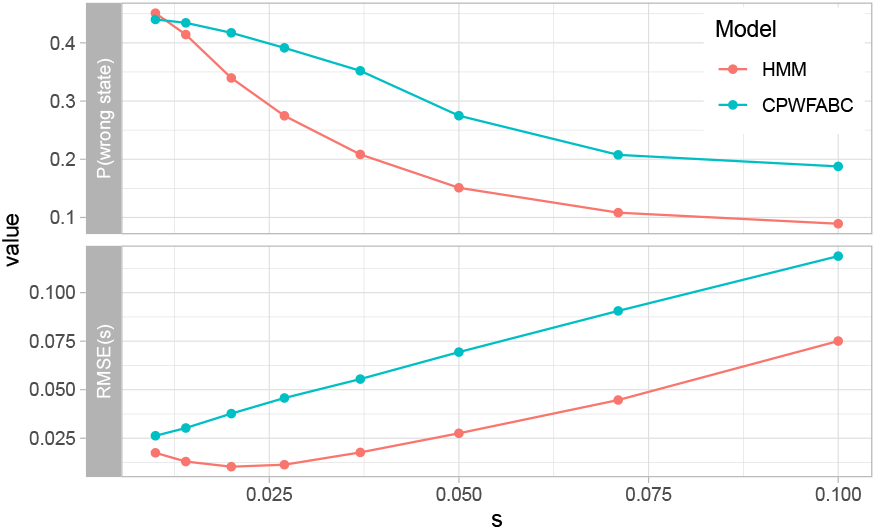
Performance comparison with CP-WFABC. We show the probability of being in the wrong state ±10 generations around the true changepoint, and the average error in the estimated selection coefficient (i.e. 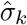 for our HMM and the CP-WFABC posterior mode).

**Figure S7:**
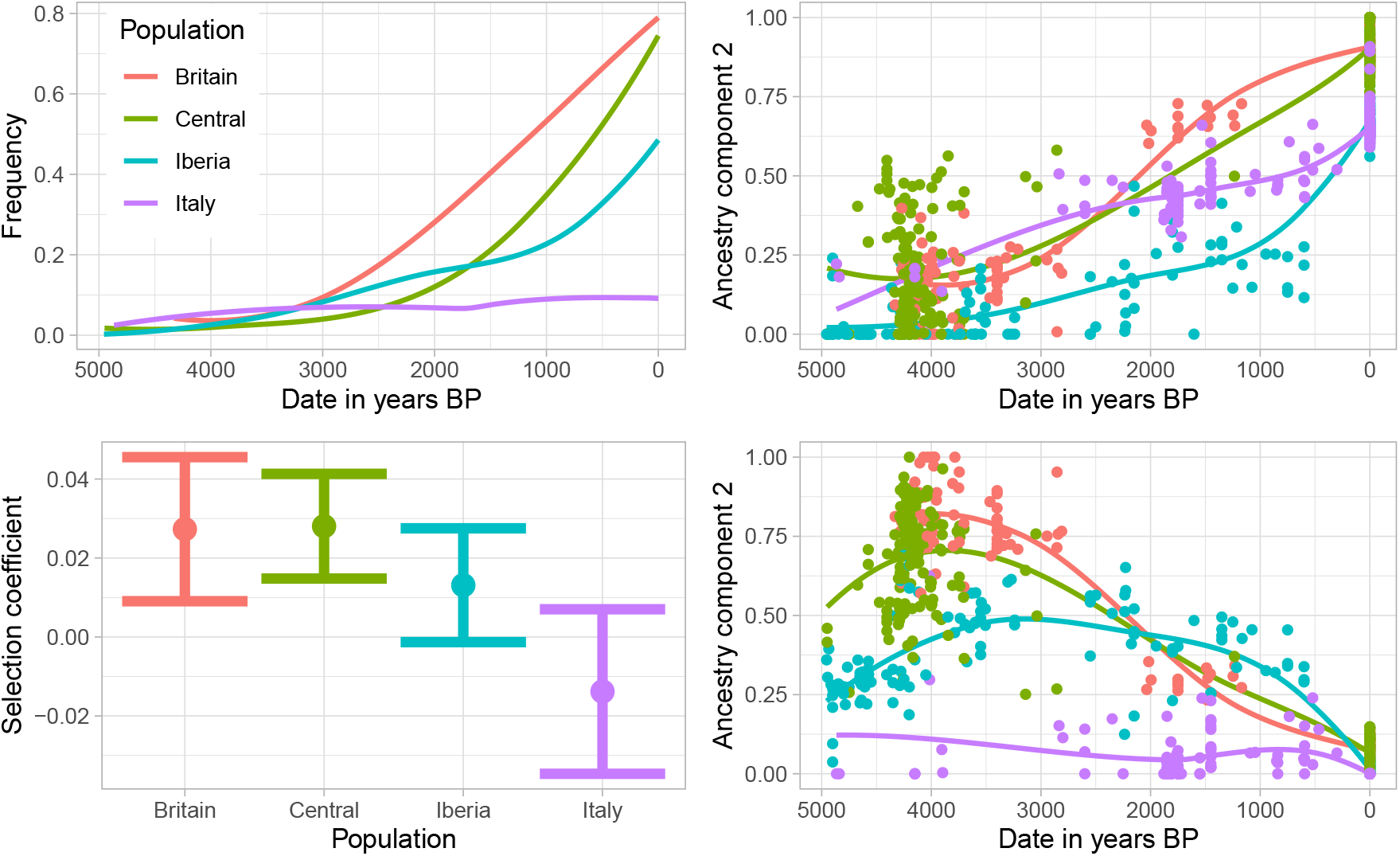
Results of fitting a logistic regression to the observations of the derived *LCT* allele, as a function of date and ancestry (inferred using ADMIXTURE with *K* = 3, and converting the effect size for date to an estimate of the selection coefficient (Methods). **Top left**: LOESS smoothed fitted allele frequency trajectories in each region. **Top left**: Estimated selection coefficients and 95% confidence intervals in each region **Right panels**: Ancestry components for each individual, with smoothed LOESS fit lines.

**Figure S8:**
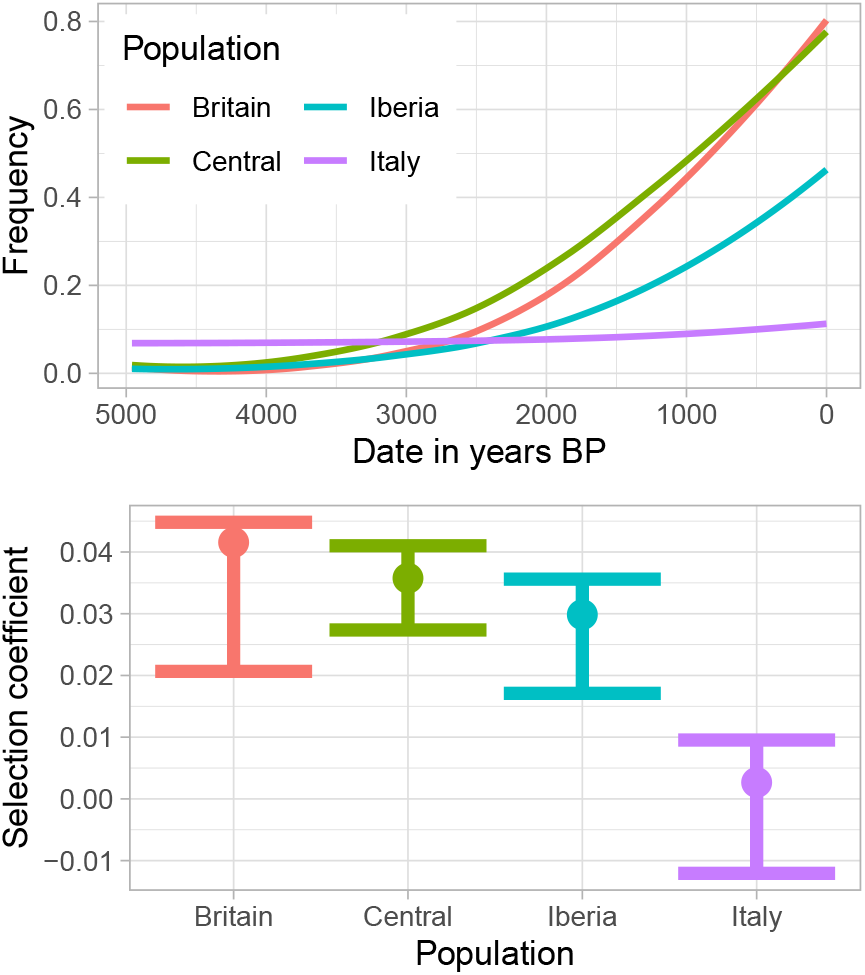
Results of fitting the 2 × 2 lattice model of (Mathieson and McVean, 2013) to the data, allowing migration between Britain and Iberia, Britain and Central, Central and Italy, and Italy and Iberia. **Upper panel**: Inferred allele frequency trajectories in each region. **Lower panel**: Estimated selection coefficients and approximate 95% confidence intervals in each region.

**Figure S9:**
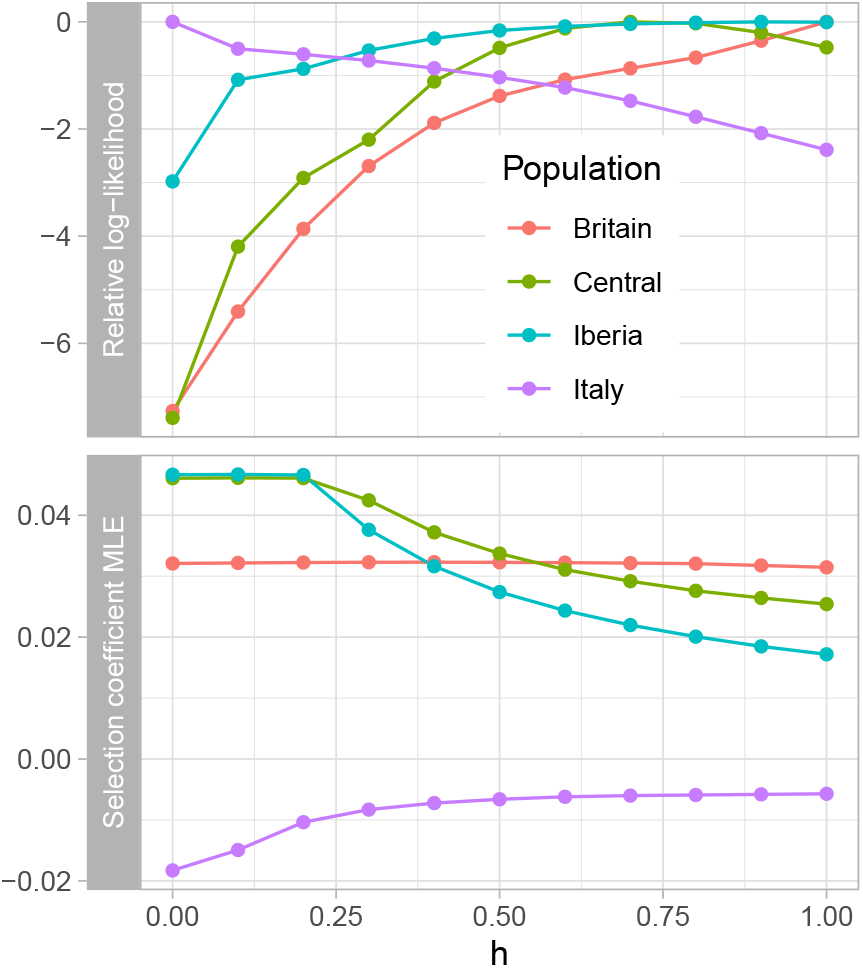
Results of fitting the single population model of Mathieson and McVean (2013) and allowing the dominance parameter *h* to vary. **Upper panel**: Log-likelihood (relative to the maximum) as a function of the dominance parameter *h*. **Lower panel**: Maximum likelihood estimate of *s* as a function of the dominance parameter *h*.

**Figure S10:**
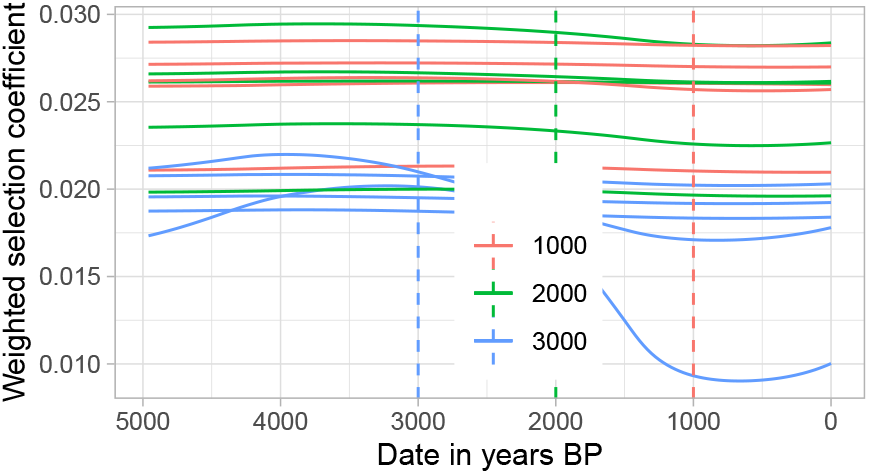
Testing whether we can detect the end of selection on *LCT*. Keeping the existing sampling points, we made the inferred allele frequency trajectory 1000, 2000 or 3000 years shorter, keeping the same total increase in frequency and inserting a 500,1000 or 1500 year period of constant frequency until the present. We then simulated observations keeping the observed distribution, and reran the estimator. We show 5 replicate simulations for each estimator.

**Figure S11:**
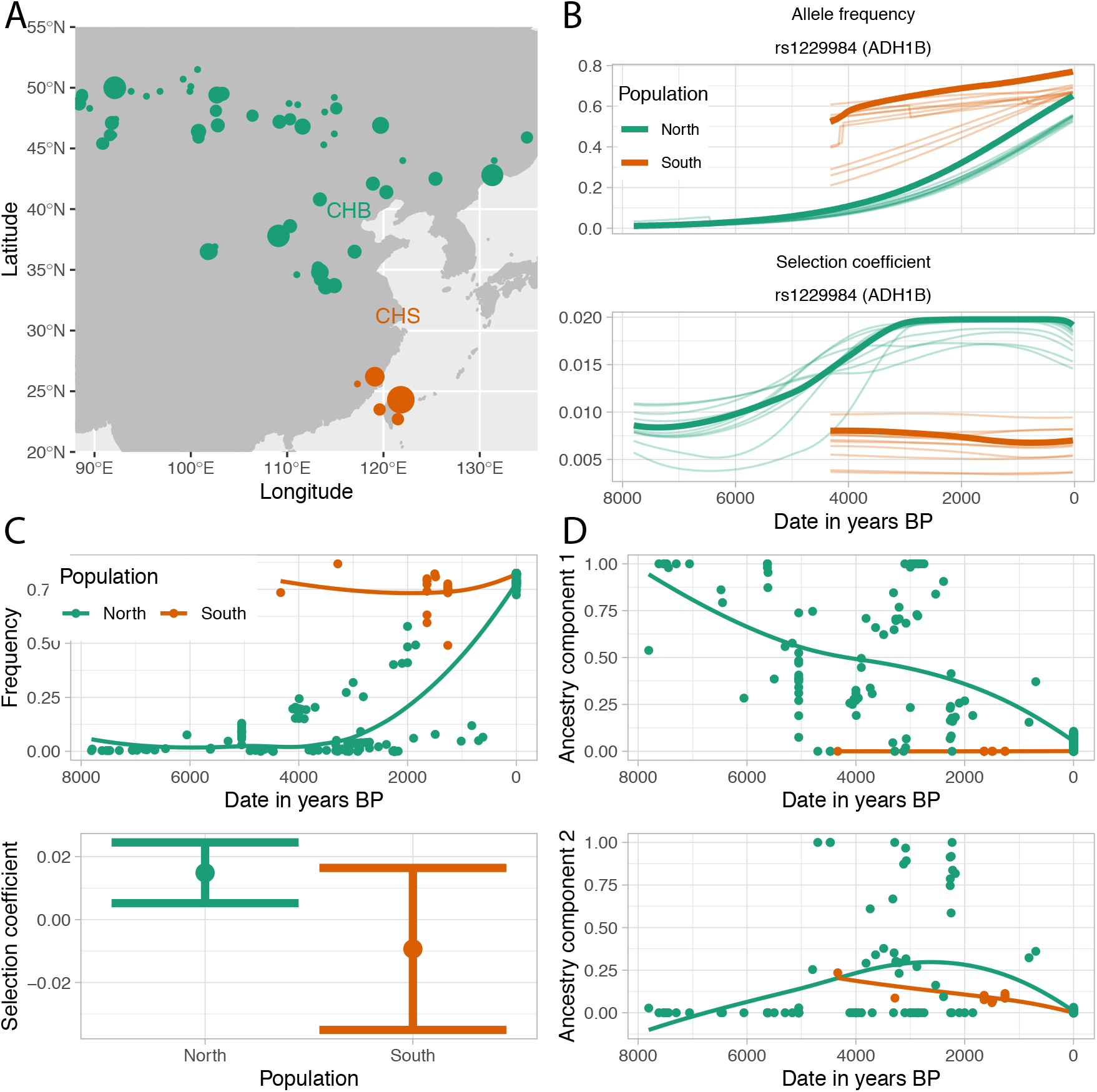
**A**: Location of present-day CHB and CHS populations from 1000 Genomes. **B**: Inferred frequency trajectories and selection coefficients for the derived ADH1B allele using present-day 1000 Genomes population frequencies (CHB/CHS). **C**: Inferred allele frequency trajectory and (constant) selection coefficient for the logistic regression model. Points show the fitted values for each ancient individuals and lines show a LOESS fit. **D**: Two ancestry components inferred using ADMIXTURE. Points show the fitted values and lines show a LOESS fit.

**Figure S12:**
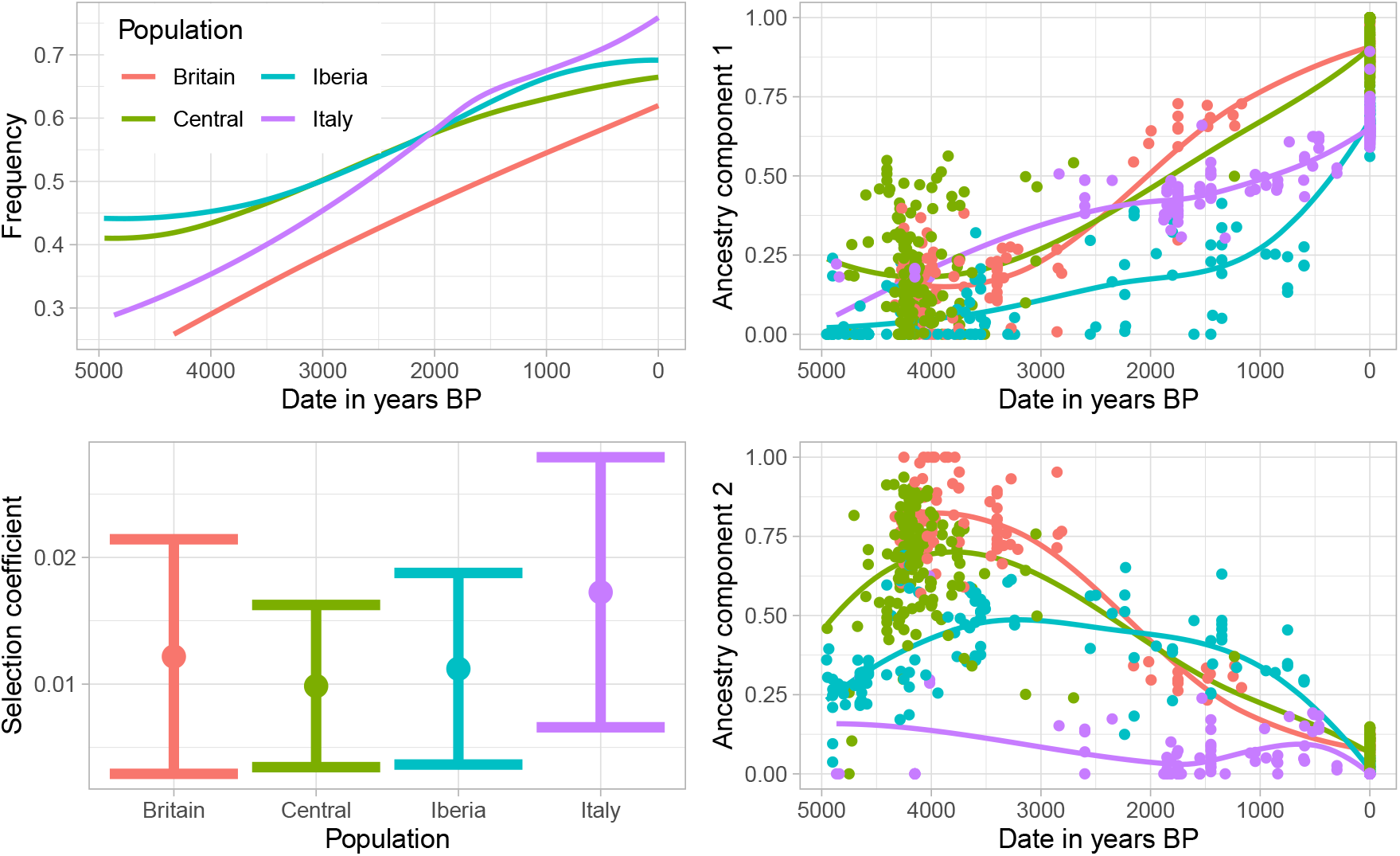
Results of fitting a logistic regression to the observations of the derived *FADS* allele in Europe, as a function of date and ancestry (inferred using ADMIXTURE with *K* = 3), and converting the effect size for date to an estimate of the selection coefficient (Methods). **Upper left**: Fitted allele frequency trajectories in each region. **Lower left**: Estimated selection coefficients and 95% confidence intervals in each region **Right panels**: Ancestry components for each individual (identical to Figure S7), with region-specific smoothed loess fit lines.

**Figure S13:**
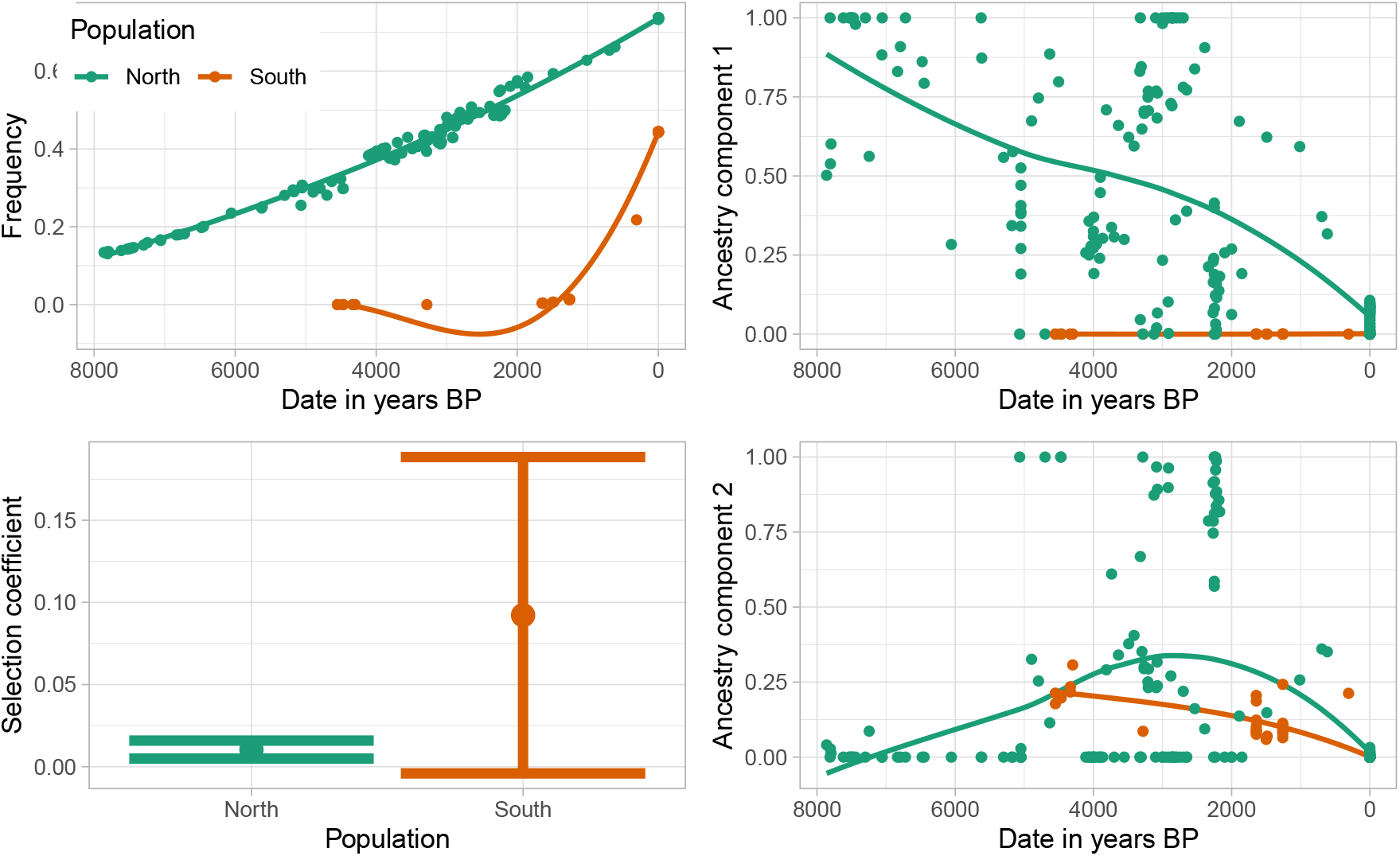
Results of fitting a logistic regression to the observations of the derived *FADS* allele in East Asia, as a function of date and ancestry (inferred using ADMIXTURE with *K* = 3), and converting the effect size for date to an estimate of the selection coefficient (Methods). **Upper left**: Fitted allele frequency trajectories in each region. **Lower left**: Estimated selection coefficients and 95% confidence intervals in each region (0.004-0.015 and −0.01-0.18 in North and South, respectively). **Right panels**: Ancestry components for each individual (identical to Figure S11), with region-specific smoothed LOESS fit lines.

